# PARP inhibition in Ewing sarcoma: impact of germline DNA damage repair defects and activation of immunoregulatory pathways

**DOI:** 10.1101/2020.09.18.304238

**Authors:** Lisa M. Maurer, Rosemarie E. Venier, Elina Mukherjee, Claire M. Julian, Jessica D. Daley, Nathanael G. Bailey, Michelle F. Jacobs, Chandan Kumar-Sinha, Haley Raphel, Kurt Weiss, Katherine A. Janeway, Rajen Mody, Peter C. Lucas, Linda M. McAllister-Lucas, Kelly M. Bailey

## Abstract

Ewing sarcoma, an oncofusion-driven primary bone tumor, can occur in the setting of various germline mutations in DNA damage repair pathway genes. We recently reported our discovery of a germline mutation in the DNA damage repair protein *BARD1* (BRCA1-associated RING domain-1) in a patient with Ewing sarcoma. BARD1 is recruited to the site of DNA double stranded breaks via the poly(ADP-ribose) polymerase (PARP) protein and plays a critical role in DNA damage response pathways including homologous recombination. PARP inhibitors (PARPi) are effective against Ewing sarcoma cells *in vitro*, though have demonstrated limited success in clinical trials to date. In order to assess the impact of BARD1 loss on Ewing sarcoma sensitivity to PARP inhibitor therapy, we generated the novel PSaRC318 patient-derived Ewing tumor cell from our patient with a germline *BARD1* mutation and then analyzed the response of these cells to PARPi. We demonstrate that PSaRC318 cells are sensitive to PARP inhibition and by testing the effect of BARD1 depletion in additional Ewing sarcoma cell lines, we confirm that loss of BARD1 enhances PARPi sensitivity. In certain malignancies, DNA damage can activate the IRF1 (interferon response factor 1) immunoregulatory pathway, and the activation of this pathway can drive immunosuppression through upregulation of the immune checkpoint protein PD-L1. In order to determine the ability of PARPi to alter Ewing tumor immunoregulation, we evaluated whether PARPi results in upregulation of the IRF1-PDL1 pathway. Indeed, we now demonstrate that PARPi leads to increased PD-L1 expression in Ewing sarcoma. Together, these data thus far suggest that while Ewing tumors harboring germline mutations in DNA damage repair proteins may in respond to PARPi *in vitro, in vivo* benefit of PARPi may only be demonstrated when counteracting the immunosuppressive effects of DNA damage by concurrently targeting immune checkpoint proteins.

## INTRODUCTION

Ewing sarcoma is a primary bone cancer driven by an aberrant fusion between *EWSR1* and a gene encoding an E26 transformation-specific (ETS) transcription factor, most commonly *FLI1* [1]. Mechanistically, EWS-FLI1 fusions are formed as the result of either reciprocal translocation or chromoplexy events [2]. Recent studies have revealed that EWS-FLI1 itself impairs homologous recombination (HR) by sequestering the HR protein BRCA1 (breast cancer gene 1) [3]. The disruption of HR by EWS-FLI1 supports the categorization of Ewing sarcoma as a ‘BRCAness’ tumor, phenotypically mimicking loss of BRCA1 expression [4]. Clinically, Ewing tumors demonstrate sensitivity to DNA damaging agents such as doxorubicin [5]. Preclinical studies using Ewing sarcoma cell lines and Ewing xenografts in immunodeficient mice have demonstrated significant tumor sensitivity to compounds that prevent DNA damage repair such as poly(ADP-ribose) polymerase 1 (PARP1) inhibitors (PARPi) [6-8]. This stands in contrast to the clinical benefit of PARPi monotherapy or PARPi plus cytotoxic chemotherapy observed in patients with Ewing sarcoma to date, which has been minimal [9, 10].

In addition to the DNA damage repair defects imparted by EWS-FLI1 itself, germline mutations in DNA damage repair genes, such as *APC, BRCA1, FANCC*, and *RAD51*, have been identified in greater than 10% of patients with Ewing sarcoma [11]. We recently contributed to this growing body of literature by reporting our discovery of a paternally inherited germline frameshift mutation in the RING domain of *BARD1* (BRCA1-associated RING domain protein 1) in a patient with Ewing sarcoma [12]. BARD1, a tumor suppressor, is an obligatory binding partner of BRCA1. The BRCA1-BARD1 heterodimer functions as an E3 ubiquitin ligase and promotes DNA double-strand break repair by homologous recombination [13]. The contribution of germline mutations in DNA damage repair genes, such as *BARD1*, to the overall sensitivity of Ewing tumors to DNA damage is largely unknown. Patients with BRCA1/2 mutant hereditary breast and ovarian cancers demonstrate significant response to PARPi/DNA damaging combinations, a response not seen in patients without these germline mutations [14, 15]. Thus, we questioned whether the subset of patients with Ewing sarcoma who also harbor germline mutations in DNA damage repair genes, such as our patient with a germline *BARD1* mutation, may demonstrate enhanced sensitivity to PARPi/DNA damaging agent combinations.

Significant cross-talk exists between DNA damage and activation of inflammatory/immune response signaling pathways [16-18]. Hereditary breast and ovarian cancers have been shown to upregulate expression of the immune checkpoint protein PD-L1 (programmed death-ligand 1) in response to DNA damage [19], thus promoting an immunosuppressive tumor microenvironment. Relatively little is known about Ewing sarcoma immunobiology, largely due to the small number of patients and the current lack of an immune competent animal model in which to study this cancer [20]. Given the spectrum of inherent defects in DNA damage repair present within Ewing tumors, we speculated that DNA-damage mediated immunoregulatory pathways may be activated following DNA damage in Ewing sarcoma.

Here, we demonstrate that additional hits to tumor DNA damage machinery, such loss of BARD1 expression, can indeed render Ewing cells more sensitive to PARPi. Interestingly, we also show that treatment of Ewing cells with PARPi results in activation of the IRF1/PD-L1 pathway. We thus postulate that PARPi-mediated immunosuppression may represent an explanation as to why PARPi has demonstrated minimal clinical benefit to date in advanced Ewing sarcoma. These findings lead us to propose that PARPi may be more effective in treating Ewing sarcoma when used in combination with agents aimed at reversing tumor immune suppression.

## MATERIALS AND METHODS

### Reagents

Antibodies were purchased from the following sources: Anti-CD99 FITC conjugated (BD Biosciences, cat no: 555688), anti-FLI1 (Abcam, cat no: 133485), anti-BARD1 (Bethyl Laboratories, cat no: A300-263A), anti-Phospho (Ser 139)-γH2A.X (Millipore Sigma, cat no: 05-636), goat anti-mouse IgG AF-488 (Thermo Fisher, cat no: A-11001), anti-PD-L1 PE conjugated (R&D Systems cat no: FAB1561P), anti-PD-L1 (Cell Signaling, clone E1L3N, cat no: 13684S), tubulin (Cell Signaling, cat no: 2144S), vinculin (Cell Signaling, clone E1E9V, cat no: 13901S), and anti-rabbit IgG-HRP (Promega, cat no: W401B). Additional specialized reagents include: BMN 673 (Talazoparib) (Cayman Chemical, cat no: 19782), MK-4827 tosylate (Niraparib) (Cayman Chemical, cat no: 20842), and DMSO (MP Biomedicals, cat no: 196055).

### Patient-derived relapsed Ewing sarcoma organoids and cell lines

The PSaRC318 patient derived Ewing sarcoma cell line was generated by our laboratory (IRB approved STUDY19030108 and the IRB approved Musculoskeletal Oncology Biobank and Tumor Registry). Briefly, fresh tumor biopsy tissue was placed in warm IMDM media supplemented with 20% heat-inactivated FBS. The tumor was then physically dissociated using a disposable stainless steel blade, and individual tumor pieces were embedded in growth factor-reduced Matrigel (Corning). The dissociated tumor was incubated at 37^°^C and 5% CO2, with media exchanged every 72 hours. Organoids grew over the next 3 weeks and were then harvested. The PSaRC318 Ewing sarcoma cell line was generated by: generating single cell suspensions, culturing cells on fibronectin-coated plates and serially depleting cultures of fibroblasts. STR profiling was performed on the PSaRC318 cell line in order to confirm identity prospectively.

### Additional cell lines and culture conditions

A673, CHLA9, CHLA10 and TC71 Ewing sarcoma cell lines were provided by Dr. Elizabeth Lawlor (University of Michigan, current location Seattle Children’s). A673 and TC71 cells were cultured in RPMI +L-glutamine media supplemented with 10% FBS. CHLA9 and CHLA10 cells were cultured in IMDM +L-glutamine media supplemented with 20% FBS and 1% insulin-selenium-transferrin (ITS, R&D Systems, cat no: AR013). Cell lines were maintained at 37^°^C and 5% CO2. All cells lines undergo routine STR profiling (University of Arizona Genetics Core, Tuscon, AZ) and regular monitoring for mycoplasma contamination.

### siRNA

BARD1 knockdown in Ewing sarcoma cells was achieved using *BARD1* SMART pool (Dharmacon, cat no: L-003873-00-0005). ON-TARGET plus non-targeting pool (Dharmacon, cat no: D-001810-10-2) was used as the control for all siRNA-based experiments. Briefly, 500 μL of optiMEM (Gibco, cat no: 11058021) was placed per well (6-well plate) along with 3 μL of 20 microMolar siRNA (reconstituted per manufacturer’s instructions using 5x siRNA buffer, Dharmacon, cat no: B-002000-UB-100) and 2 μL (A673) or 2.5 μL (CHLA10) Liopfectamine RNAiMAX transfection reagent (Life Technologies, cat no: 13778150). The mixture was incubated with intermittent rocking for 30 minutes prior to the addition of cells.

### Radiation

Cells were radiated with X-RAD 320 (Precision X-ray Inc, N. Branford, CT USA) using Filter 2 (1.5mm Al + 0.25mm Cu +0.75mm Sn) at doses (Gy) noted in individual experiments.

### RT-PCR

RNA isolation was performed using Qiagen RNeasy Plus isolation kit (Qiagen, cat no: 74134) for >500K cells or Qiagen RNeasy Plus Micro isolation kit (Qiagen, cat no: 74034) for ≤500K cells. RNA concentration was measured using Nanodrop (Thermo Scientific, Waltham, MA, USA). cDNA synthesis was performed on 1 µg of RNA with the high-capacity cDNA reverse transcription kit (ThermoFisher Scientific, cat no: 4374966) and Applied Biosystems Veriti 96-well thermocycler.

qRT-PCR analysis was performed using Taqman probes (Life Technologies, GAPDH Hs02758991_g1, RPLP0 Hs00420895_gH, BARD1 Hs00957655_m1, PD-L1 (CD274) Hs00204257_m1 and IRF1 Hs00971965), Taqman Universal PCR Master Mix (Life Technologies, cat no: 4304437), and StepOnePlus Real-Time PCR system (Life Technologies).

### Immunoblotting

Immunoblot analysis was performed on cell lysates prepared using LDS lysis buffer as previously reported [21]. After sonication, gel electrophoresis was performed using SDS-PAGE gel electrophoresis. After transfer to nitrocellulose, membranes were blocked with 5% milk in TBST and incubated with primary antibody overnight in 2.5-5% milk TBST at 4°C. After washing, membranes were incubated with HRP conjugated secondary antibody prior to the addition of ECL reagent (Thermo Scientific).

### Flow cytometry

Adherent Ewing sarcoma cells were detached from the culture plate using Accutase Cell Detachment Solution (Corning, cat no: 25-058-Cl). Cells were first stained with Live/Dead Aqua (Life Technologies, #L34957) using a ratio of 1 μL stain per 1×10^6 cells in 1mL. Cells were then stained for either CD99 FITC (BD Biosciences, cat no: 555688) using 5 μL of antibody per 1×10^6 cells in 1mL or PD-L1 PE using 10 μL of antibody per 1×10^6 cells in a total volume of 100μL. The percentage of live, FITC or PE positive cells and the mean fluorescence intensity was determined using a BD Aria IIu or BD Aria II SORP. FloJo software was used for data analysis and generation of data plots.

### RNA-seq

The University of Pittsburgh Health Sciences Sequencing Core at the UPMC Children’s Hospital of Pittsburgh performed RNA extraction from FFPE tumor tissue, measured RNA quantity and quality using Qbit (Thermo Fisher Scientific), and performed mRNA library preparation and RNA-Seq using an Illumina platform. R version 4.02 and package fgsea version 1.14.0 were used for data analysis and figure generation.

### Immunofluoresence staining and confocal microscopy

Cells were fixed and immunofluorescently labeled as previously described [22]. Slides were mounted using a DAPI-containing mounting solution and cells were imaged using a Zeiss LSM 510 confocal microscope.

### Live-cell monitoring and apoptosis assays

Cells were seeded into 96 well plates (Corning 3610 or 3596) in 100 μL of Fluorobrite media (Gibco, #A18967-01) containing 5% FBS. Cell treatment conditions are described in the individual experiments. For apoptosis assays, IncuCyte Caspase 3/7 green reagent (Essen BioScience, cat no: 440) was added to a final dilution of 1:1000. Phase contrast images of the cells in standard culture conditions were obtained at 3-6 hour intervals using an IncuCyte S3 or IncuCyte Zoom (Essen BioScience). Green fluorescence images were additionally captured for apoptosis assays.

### Cancer Cell Line Encyclopedia (CCLE) analysis

Ewing sarcoma cell lines were queried in the Broad Institute Cancer Cell Line Encyclopedia (CCLE) database (https://portals.broadinstitute.org/ccle). The mutations in each cell line were compared against genes involved in DNA damage repair and classified as pathogenic, likely pathogenic, variant of unknown significance, benign or likely benign using COSMIC.

### PEDS MiONCOseq data analysis

PEDS-MiONCOseq is an IRB-approved, pediatric precision oncology pediatric cohort enrolling since May 2011 as previously described [23]. The whole exome sequencing data from this cohort of now >500 pediatric oncology patients was queried for germline mutations in *BARD1*. Pathogenicity status of mutations identified was determined.

### St. Jude Could PeCan data analysis

The St. Jude Cloud PeCan (Pediatric Cancer Knowledgebase) (https://www.stjude.cloud, doi:https://doi.org/10.1101/2020.08.24.264614) germline sequencing data was queried for germline mutations in *BARD1* and corresponding pathogenicity status of mutations.

### Statistical analyses

PRISM software was used in order to plot individual data points and the standard deviation. Single comparisons were performed through the use of an unpaired, two-tailed Student *t* test.

## RESULTS

### The landscape of germline *BARD1* mutations in pediatric oncology patients

Pathogenic germline *BARD1* mutations provide a moderate risk for heritable breast cancer and have also been reported in pediatric patients diagnosed with high risk neuroblastoma [24-26]. We previously reported our discovery of a pathogenic germline *BARD1* mutation in a patient with Ewing sarcoma [12]. To better understand the frequency of germline *BARD1* mutations in pediatric malignancies, we investigated the presence of germline *BARD1* mutations in two pediatric oncology sequencing databases: PEDS-MiONCOseq and St. Jude Cloud PeCan (**Fig. 1**). Pathogenic or likely pathogenic (red) germline *BARD1* mutations were found in patients diagnosed with multiple different cancers including neuroblastoma, glioma, and B-cell acute lymphoblastic leukemia. Three patients with bone sarcomas were identified as having germline *BARD1* variants of unknown significance (VUS, denoted in blue). The EWS-FLI1 fusion oncoprotein can bind to BARD1 [27] and it is unknown how the presence of germline or somatic *BARD1* VUS may alter EWS-FLI1 interactions with the protein. Benign or likely benign germline *BARD1* mutations were also noted in this pediatric cancer patient cohort and are included in **Supplemental Figure 1**.

**Figure 1.**
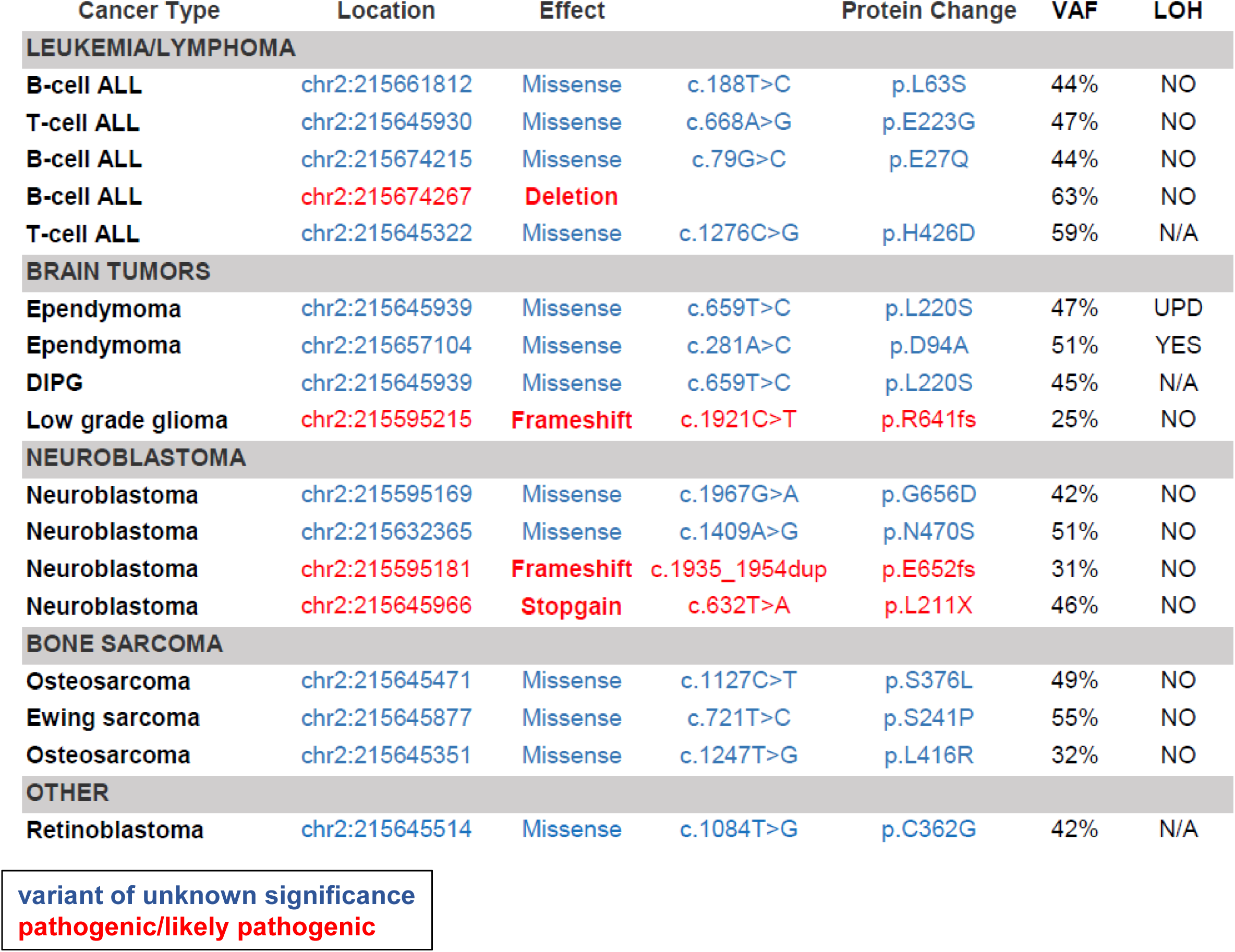
The landscape of germline *BARD1* mutations in a cohort of pediatric oncology patients. Germline sequencing data from PEDS MiONCOseq and St. Jude PeCan datasets was queried for mutations in *BARD1* and the pathogenicity status of the individual mutations were determined. Mutations highlighted in blue are variants of unknown significance. Mutations highlighted in red are pathogenic or likely pathogenic. Abbreviations: ALL=Acute Lymphoblastic Leukemia; LOH=loss of heterozygosity, VAF=variant allele frequency, UPD=uniparental disomy, and N/A=data not available.

### PSaRC318: a patient-derived Ewing sarcoma cell line

Germline DNA damage repair mutations are reported in >10% of patients with Ewing sarcoma [11]. We have previously reported a patient with Ewing sarcoma who harbors a pathogenic germline *BARD1* mutation that results in a frameshift that introduces a premature stop codon within the RING domain of BARD1 [12]. This patient was enrolled in our IRB-approved Musculoskeletal Oncology Biobank and Tumor Registry (MOTR) and viable tumor was banked at the time of relapse. RNA-seq analysis performed on tumor obtained at the time of relapse demonstrated an EWS-FLI1 signature by gene set enrichment analysis (**Fig. 2A**). Next, viably frozen tumor tissue from biopsy of the relapsed lung lesion was used to develop a novel Ewing sarcoma cell line, PSaRC318, the morphology of which is shown in **Fig. 2B**. Flow cytometry confirmed that greater than 99% of PSaRC318 cells express surface CD99, a commonly used marker to identify Ewing sarcoma cells (**Fig. 2C**). Sequencing from the tumor revealed a rare Type 3 EWS-FLI1 fusion (data not shown). Type 3 fusions contain a larger N-terminal portion of EWS (exon 10 breakpoint) as compared to the more common Type 1 EWS-FLI1 fusion (exon 7 breakpoint) (see **Fig. 2D**, schematic). Expression of the Type 3 fusion oncoprotein was confirmed via anti-FLI1 Western blot analysis of PSaRC318 lysates (**Fig. 2D**). In comparison with other Ewing sarcoma cell lines (A673, TC71, CHLA9, and CHLA10), PSaRC318 cells demonstrate significantly reduced expression of BARD1 (**Fig. 2E**).

**Figure 2.**
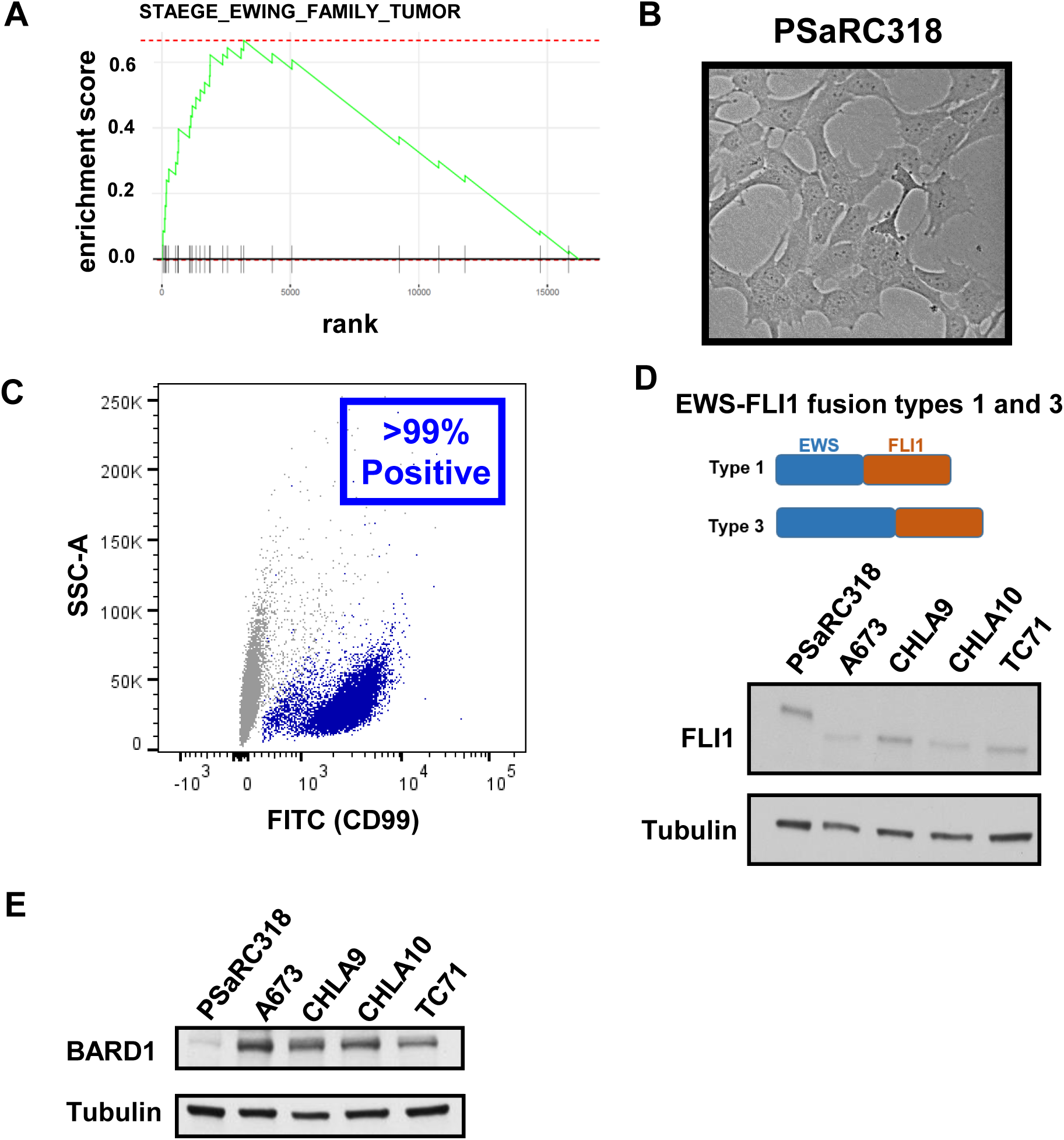
Characterization of PSaRC318, a primary Ewing tumor cell line from a patient with a *BARD1* frameshift mutation. **A**, RNAseq analysis performed on relapsed tumor demonstrates enrichment of the STAEGE_EWING_FAMILY_TUMOR geneset upon geneset enrichment analysis (GSEA). **B**, Phase contrast image of the PSaRC318 Ewing tumor cell line. **C**, Flow cytometry showing presence of surface CD99 expression in the PSaRC318 cell line. **D**, Schematic detailing the difference between Type 1 and Type 3 EWS-FLI fusions (top) and Western blot with anti-FLI1 antibody of Ewing sarcoma cell lines with type 3 (PSaRC318) versus type 1 (A673, CHLA9, CHLA10, and TC71) EWS-FL1 fusions. **E**, Western blot demonstrating BARD1 protein expression in the same Ewing sarcoma cell lines as in (**D**).

### Loss of BARD1 enhances Ewing tumor cell sensitivity to PARP inhibition

Next, we questioned whether PSaRC318 cells would be sensitive to PARPi. To verify that talazoparib induced DNA damage in our model system, PSaRC318 cells were treated with talazoparib (versus DMSO control) for 48 hours, fixed, and then labeled for phospho-γH2AX (p-γH2AX), a marker of DNA double strand breaks (**Fig. 3A**). PSaRC318 indeed showed an increase in p-γH2AX staining after treatment with the talazoparib as compared with DMSO control. To determine the impact of talazoparib treatment on the growth and survival of PSaRC318 cells, IncuCyte assays were performed to monitor PSaRC318 cell growth following treatment with DMSO versus 100 nM talazoparib. We found that PSaRC318 cell growth was significantly impaired by talazoparib treatment (**Fig. 3B**). Next, in order to more directly determine the impact of BARD1 loss on Ewing cell sensitivity to PARP inhibition, we evaluated the effect of siRNA-mediated knockdown of BARD1 in A673 and CHLA10 cells. Efficient knockdown of BARD1 was achieved in both A673 and CHLA10 cells as demonstrated by qRT-PCR and Western blot analysis (**Fig. 3C, D**). Additionally, A673 and CHLA10 cell treatment with talazoparib for 24 hours enhanced punctate p-γH2AX staining (**Fig. 3E**). As compared with cells treated with control siRNA (Ctsi), A673 and CHLA10 cells treated with BARD1siRNA were both more sensitive to talazoparib in Incucyte assays (**Fig. 3F**). Together, these results strongly suggest that loss of BARD1 enhances Ewing cell sensitivity to PARP inhibition.

**Figure 3.**
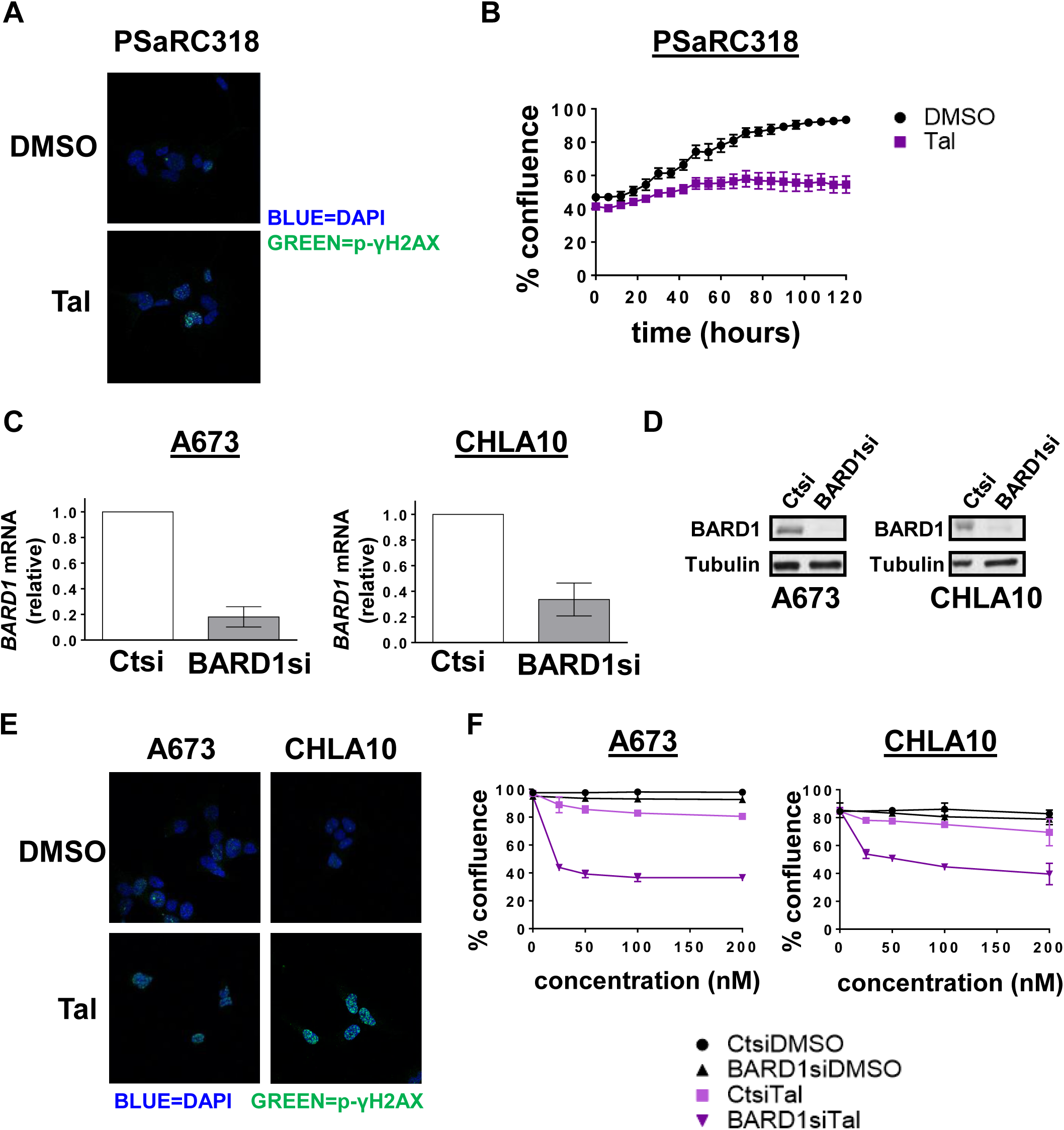
Loss of BARD1 enhances Ewing sarcoma cell sensitivity to PARP inhibition. **A**, p-γH2AX and DAPI staining of PSaRC318 cells treated with 100 nM talazoparib (Tal). **B**, IncuCyte assay comparing the growth of PSaRC318 cells treated with DMSO versus 100 nM Tal. **C**, qRT-PCR showing *BARD1* mRNA expression in A673 or CHLA10 cells treated with control (Ctsi) or BARD1 (BARD1si) siRNA. **D**, Western blot analysis showing BARD1 expression in A673 or CHLA10 cells treated with Ctsi or BARD1si. **E**, p-γH2AX and DAPI staining of A673 and CHLA10 cells treated with 100 nM Tal. **F**, IncuCyte monitoring of cell confluence at increasing concentrations of talazoparib (Tal) versus DMSO controls in A673 and CHLA10 Ewing sarcoma cells treated with Ctsi versus BARDsi. A673 and CHL10 cell data is graphed at the 60hrs time point.

### Mutations in DNA damage repair genes in Ewing sarcoma cell lines

Prior *in vitro* studies have shown that Ewing sarcoma cells are among the cancer cell lines that are most sensitive cells to PARP inhibition [6, 7]. Given our result demonstrating that mutations in DNA damage repair genes, such as *BARD1*, can significantly enhance sensitivity to PARPi, we questioned whether the Ewing cell lines used in these original studies may demonstrate a significant number of cell lines with DNA damage repair defects. To answer this question, sequencing data available for these Ewing cell lines through the Cancer Cell Line Encyclopedia (CCLE, Broad Institute) was analyzed for mutations in DNA damage repair genes. Pathogenic mutations in *FANCM* were noted in two of the Ewing cell lines. Additionally, a pathogenic *SLX4* mutation was noted in one cell line (**Fig. 4**).

**Figure 4.**
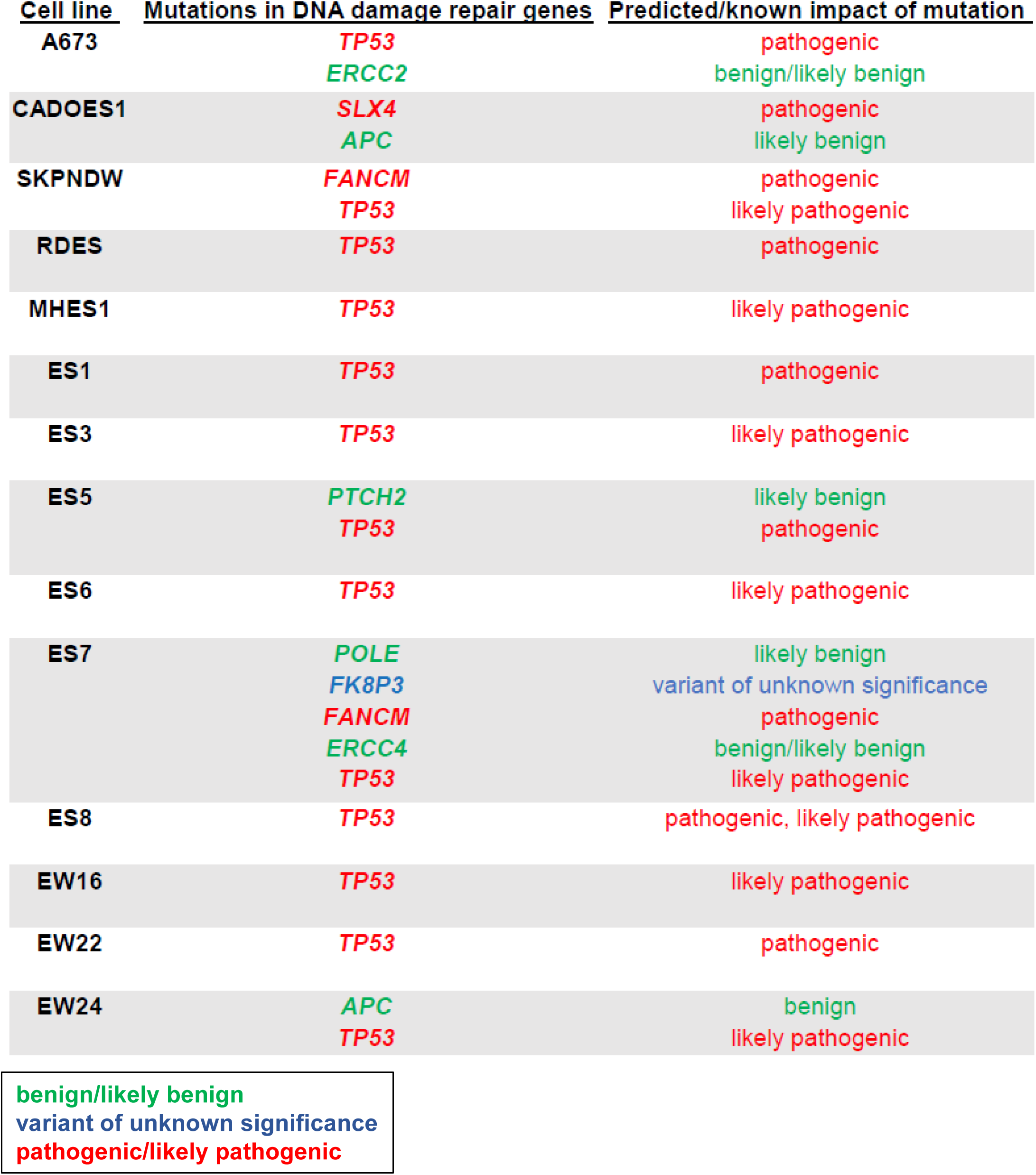
Prevalence of DNA damage repair gene mutations in Ewing cell lines included in original studies of PARP inhibition. Cancer cell line encyclopedia (CCLE) data was queried for mutations in DNA damage repair gene mutations in the available 14 Ewing sarcoma cell lines listed. The predicted/known impacts of the mutations are also noted.

### Loss of BARD1 enhances apoptosis in Ewing cells treated with PARPi plus radiation

PARPi monotherapy tends to reduce proliferation but has less of an impact on Ewing cell apoptosis. The addition of DNA damage in the setting of PARPi has been shown to enhance apoptosis of Ewing cells [28]. We next wanted to determine the impact of reducing BARD1 expression on Ewing cell response to a combination of DNA damage (radiation) plus PARPi. We performed this analysis using a second PARPi (niraparib) given that individual PARP inhibitors are known to have off target effects including the inhibition of various cellular kinases [29]. PSaRC318 cells were treated with 0.5 μM niraparib or DMSO control and then treated with or without 2Gy radiation at 15 hours and monitored via IncuCyte. PSaRC318 cells demonstrate significant apoptosis and decreased confluence in the presence of PARPi + radiation (**Fig. 5A, B)**. To more precisely examine the role of BARD1, A673 and CHLA10 cells treated with Ctsi or BARD1siRNA and then analyzed. A673 cells were treated with 0.5 or 1 μM niraparib and CHLA10 cells were treated with 1 or 1.5 μM niraparib and monitored over time using an IncuCyte. At ∼12 hours, cells were treated with or without 2 Gy radiation. In A673 cells, knockdown of BARD1 led to an increase in apoptosis and a corresponding decrease in cell confluence of cells treated with niraparib as compared with control cells treated with niraparib (**Fig. 5C, D**). Similar results were seen with CHLA10 cells (**Supplemental Fig. 2**), though as noted, higher doses of niraparib were required to achieve this effect. Together, these data demonstrate that reduction of BARD1 in Ewing sarcoma: 1) enhances sensitivity to both talazoparib and niraparib, 2) leads to increased apoptosis in the setting of PARPi plus direct DNA damage (radiation), and 3) can partially resensitize resistant Ewing cells (CHLA10) to PARPi. Based on this, we postulated that perhaps patients with Ewing sarcoma and additional germline mutations in DNA damage repair genes could represent a unique subset of patients that would demonstrate significant clinical benefit from PARPi combinatorial therapy.

**Figure 5.**
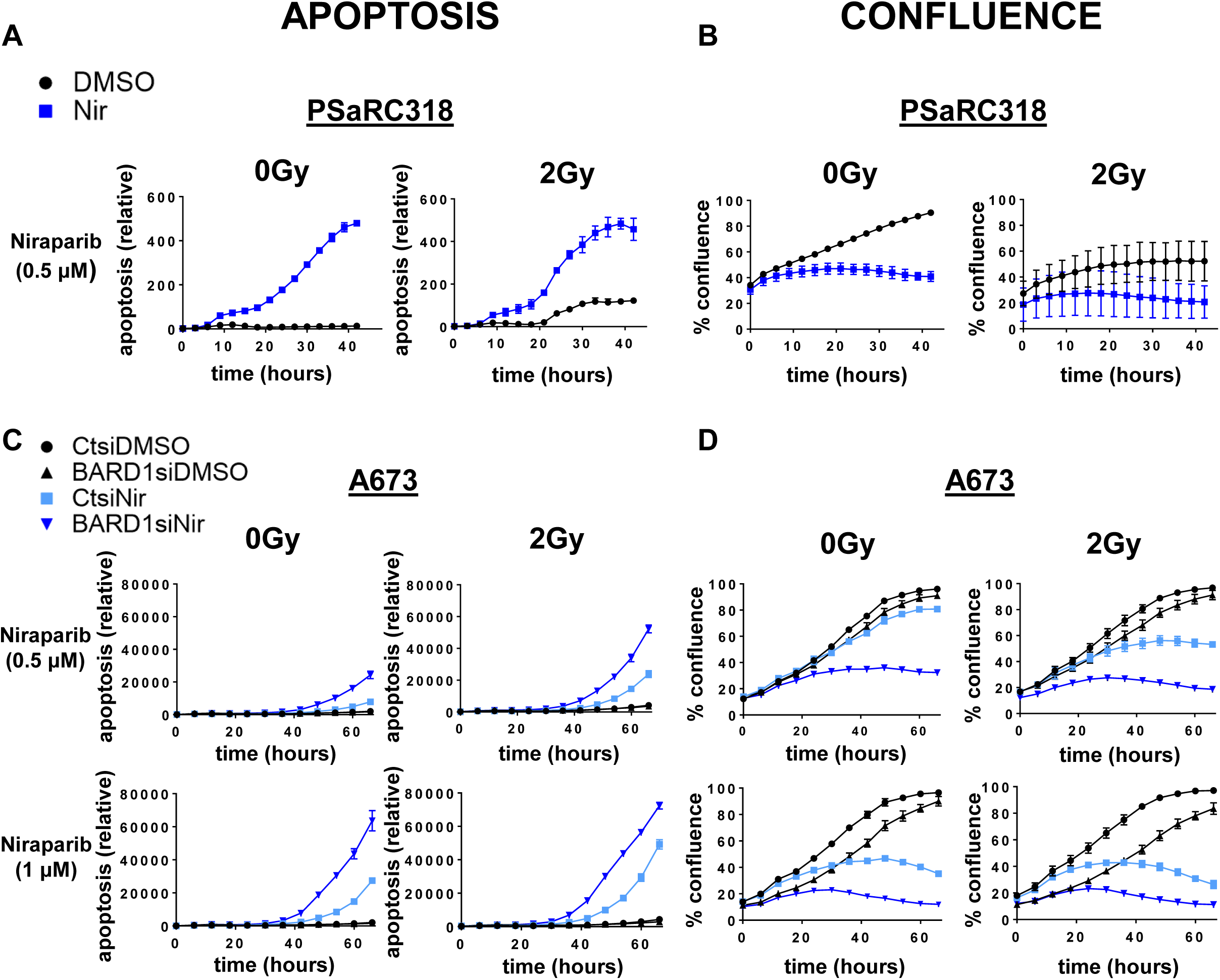
BARD1 loss enhances Ewing sarcoma cell apoptosis in response to niraparib plus radiation. **A**, Relative apoptosis data from IncuCyte assays showing the effect of 0.5 μM niraparib (Nir) versus DMSO control plus either 0 or 2 Gy radiation on PSaRC318 cells. **B**, Confluence data from IncuCyte assays showing the effect of 0.5 μM Nir versus DMSO control plus either 0 or 2 Gy radiation on PSaRC318 cells. **C, D**, A673 cells were treated with control (Ctsi) or BARD1 (BARD1si) siRNA, Nir (at doses indicated) versus DMSO control, and either 0 or 2 Gy radiation and monitored via IncuCyte apoptosis assay **(C)** or confluence assay **(D)**. For these experiments, cells were seeded in the presence of Nir and radiation was performed at 12-15 hours. Relative apoptosis (caspase 3/7 activity) is calculated as green florescence in μM^2^ divided by percent confluence.

### Ewing cells upregulate the IRF1 immunoregulatory pathway in response to PARPi

This postulated clinical benefit of PARPi in a subset of Ewing sarcoma patients could be negated by the effect of DNA damage on tumor immunoregulation. Tumors with accumulating DNA damage can dampen the immune response through activation of immunoregulatory pathways such as the IRF1 pathway. The downstream effect of IRF1 pathway activation is the upregulation of immune checkpoint proteins, such as PD-L1 [17, 30]. Thus, while DNA damage may result in apoptosis of some tumor cells, surviving cells may gain the capability to enhance their survival through the induction of immunosuppression.

To determine the effect of PARPi on the IRF1 immunoregulatory pathway in Ewing sarcoma, PSaRC318 cells were incubated with talazoparib and then analyzed for IRF1 pathway activation. Treatment with talazoparib for 48 hours resulted in a ∼2-fold increase in *IRF1* mRNA (**Fig. 6A**) and a 4-fold increase in *CD274* (mRNA encoding PD-L1) (**Fig. 6B**). A dose dependent increase in *IRF1* and *CD274* (PD-L1) was also seen in PSaRC318 cells treated with niraparib (**Figs. 6C, D**). Additionally, Western blot analysis of A673 Ewing cells demonstrates an increase in PD-L1 protein expression after 42 hours of niraparib treatment (**Fig. 6E**). Lastly, flow cytometry was performed in niraparib treated A673 cells pre-treated with Ctsi or BARD1si in order to examine surface PD-L1 expression. A673 cells treated with BARD1si demonstrated a ∼1.5-fold increase in surface PD-L1 expression after niraparib treatment as compared to DMSO control (**Fig. 6F, G**). These results suggest that PARPi can increase immunomodulatory pathways that could serve to protect Ewing sarcoma cells from immune surveillance.

**Figure 6.**
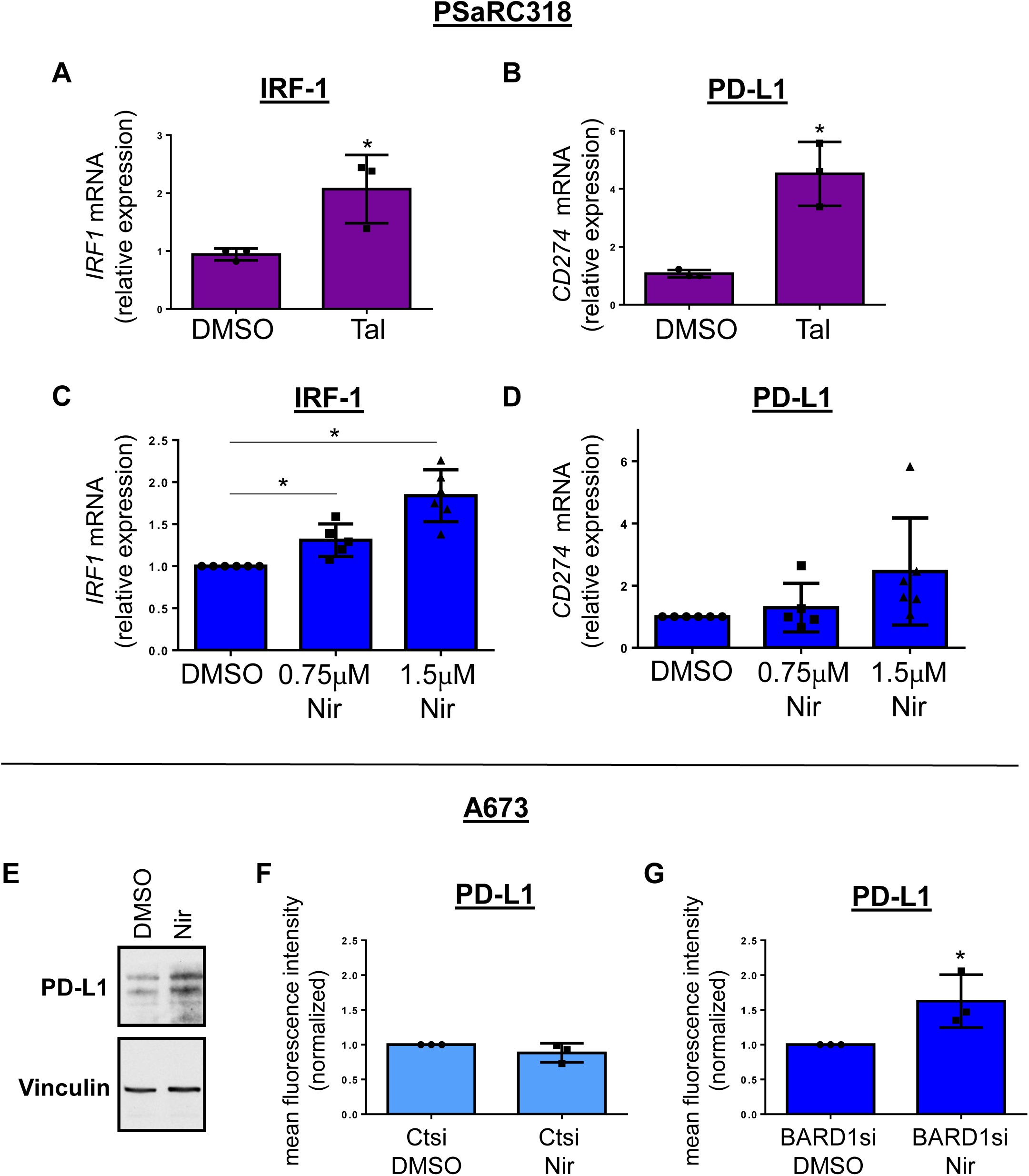
The IRF1 immunoregulatory pathway is enhanced following PARPi treatment in Ewing sarcoma. **A**, *IRF1* and **B**, *CD274*(PD-L1) mRNA expression in PSaRC318 cells treated with DMSO or talazoparib (Tal) measured using qRT-PCR. **C**, *IRF1* and **D**, *CD274*(PD-L1) mRNA expression in PSaRC318 cells treated with DMSO or niraparib (Nir) measured using qRT-PCR. **E**, Western blot of PD-L1 protein expression in A673 treated with DMSO or 1 μM Nir. Flow cytometry of PD-L1 surface expression in A673 cells pretreated with **F**, Ctsi or **G**, BARD1si and then treated with DMSO or 1 μM Nir. Mean fluorescence intensity is normalized to cells treated with DMSO. * denotes p value <0.05 calculated using an unpaired, two-tailed Student *t*test.

## DISCUSSION

Advanced Ewing sarcoma remains a deadly disease despite numerous preclinical studies suggesting the benefit of various agents, including PARP inhibitors [9, 10]. In the current work, we have generated and utilized the novel patient-derived *BARD1* mutated Ewing sarcoma cell line PSaRC318 as a model to ask the question: Can additional germline mutations in DNA damage repair genes enhance sensitivity to PARPi beyond the vulnerability already imparted by the presence of EWS-FLI1? Indeed, here we show that disrupting BARD1 expression enhances sensitivity to PARPi and enhances PARPi plus radiation-induced Ewing cell apoptosis.

While any single germline DNA damage repair gene mutation in patients with Ewing sarcoma is rare, a fact highlighted by our analysis of the PEDS MiONCOseq and St. Jude PeCan datasets, perhaps collectively, this subset of patients with germline DNA damage repair gene mutations could demonstrate greater benefit from combinatorial therapies targeting DNA damage repair machinery than patients without such mutations. Future studies will address this ongoing area of interest in our laboratory, with the goal of developing a precision medicine-based approach to optimizing treatment for patients with advanced Ewing sarcoma.

Despite the broad spectrum of DNA repair defects in Ewing tumors, PARPi have demonstrated little survival benefit in clinical trials to date [9, 10]. Upon querying the genetic composition of cell lines originally used to determine Ewing cell sensitivity to PARPi, we noted that three of the fourteen cell lines did indeed harbor additional pathogenic mutations in DNA damage repair genes. It is possible that the overrepresentation of rare DNA damage repair defective tumors in the cell lines tested could explain why PARPi have demonstrate little clinical benefit in more broad studies of patients with relapsed Ewing sarcoma.

A second factor that could negate PARPi effectiveness *in vivo* is the activation of immunoregulatory pathways following induction of DNA damage in Ewing tumors. Our current work demonstrates the upregulation of both IRF1 and PD-L1 following PARPi treatment of Ewing sarcoma cells. The upregulation of the checkpoint protein PD-L1 promotes an immunosuppressive tumor microenvironment, thus sheltering tumor cells from immune surveillance [31, 32]. Our work suggests a possible mechanism whereby Ewing cells surviving PARPi-induced DNA damage can be reprogrammed to survive through the immune-protection imparted by upregulation of the IRF1 pathway. In ovarian cancer, PARPi/checkpoint inhibitor combinations are currently being investigated as a means by which to overcome the increase in PD-L1 expression resulting from PARPi [33]. Future studies are aimed at determining the efficacy of DNA damaging agent/immunotherapy combinations in Ewing sarcoma, particularly for those patients who harbor germline loss-of-function mutations in important DNA damage repair genes such as *BRCA1* or *BARD1*.

Our work highlights the critical need to better understand both the impact of: 1) germline DNA damage repair mutations on Ewing tumor response to therapy and 2) DNA damaging agents on the Ewing tumor immune microenvironment. Currently, *in vivo* studies addressing these points are challenging given the broad lack of an immuno-competent (syngeneic or transgenic) Ewing sarcoma animal model [20]. Ongoing studies utilizing optimized *in vitro* approaches to address these points are underway.

## Acknowledgements

The authors would like to thank The Alex’s Lemonade Stand Foundation Childhood Cancer Data Laboratory’s R-learning workshop, including Dr. Huajin Wang.

## Conflict of Interest Statement

The authors have no conflicts of interest to report.

---

**Supplemental Figure 1.**
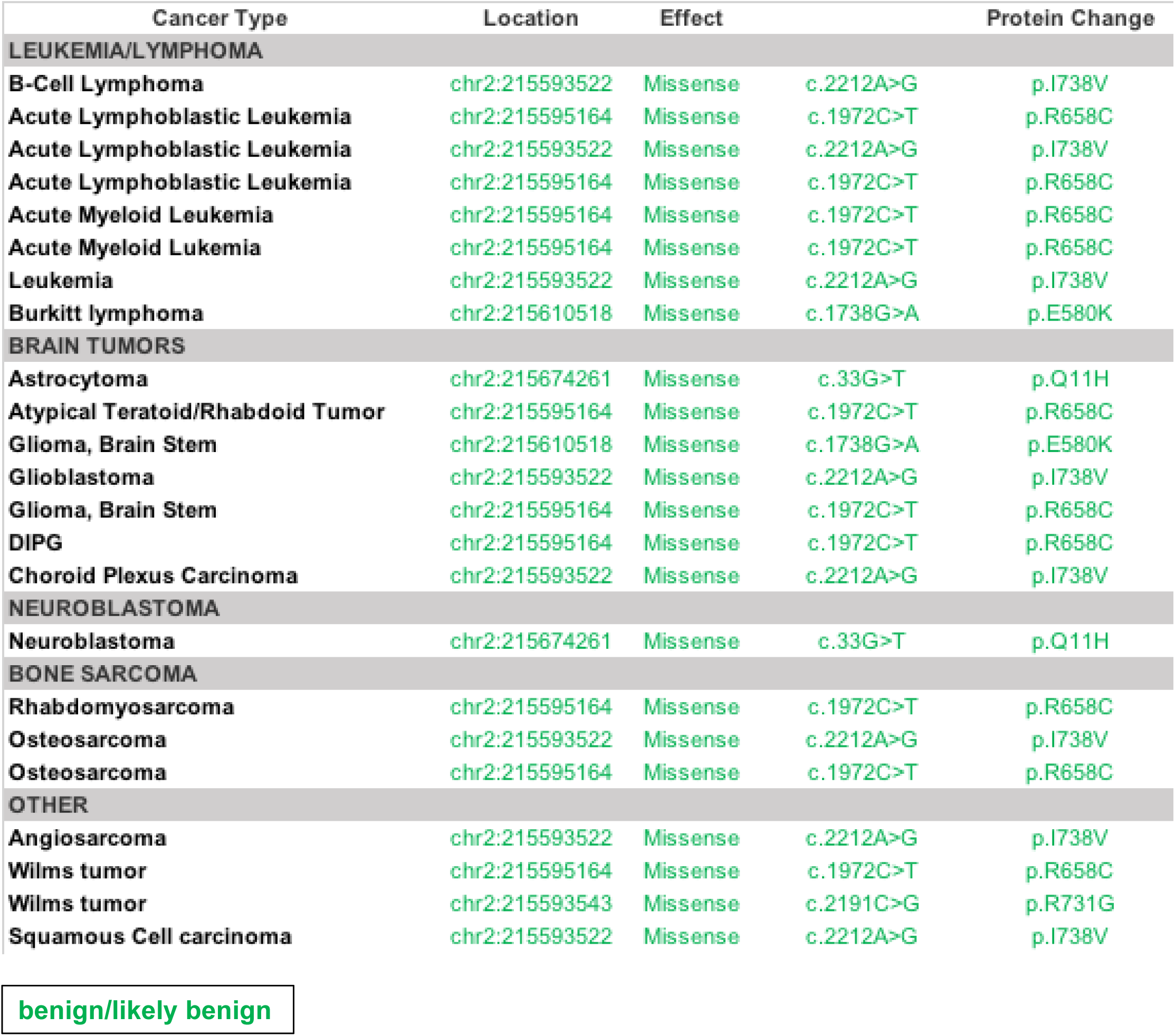
Benign germline *BARD1* mutations. Benign or likely benign germline *BARD1* mutations identified in the PEDS-MiONCOseq dataset. This serves as an extension of the data presented in **Figure 1**.

**Supplemental Figure 2.**
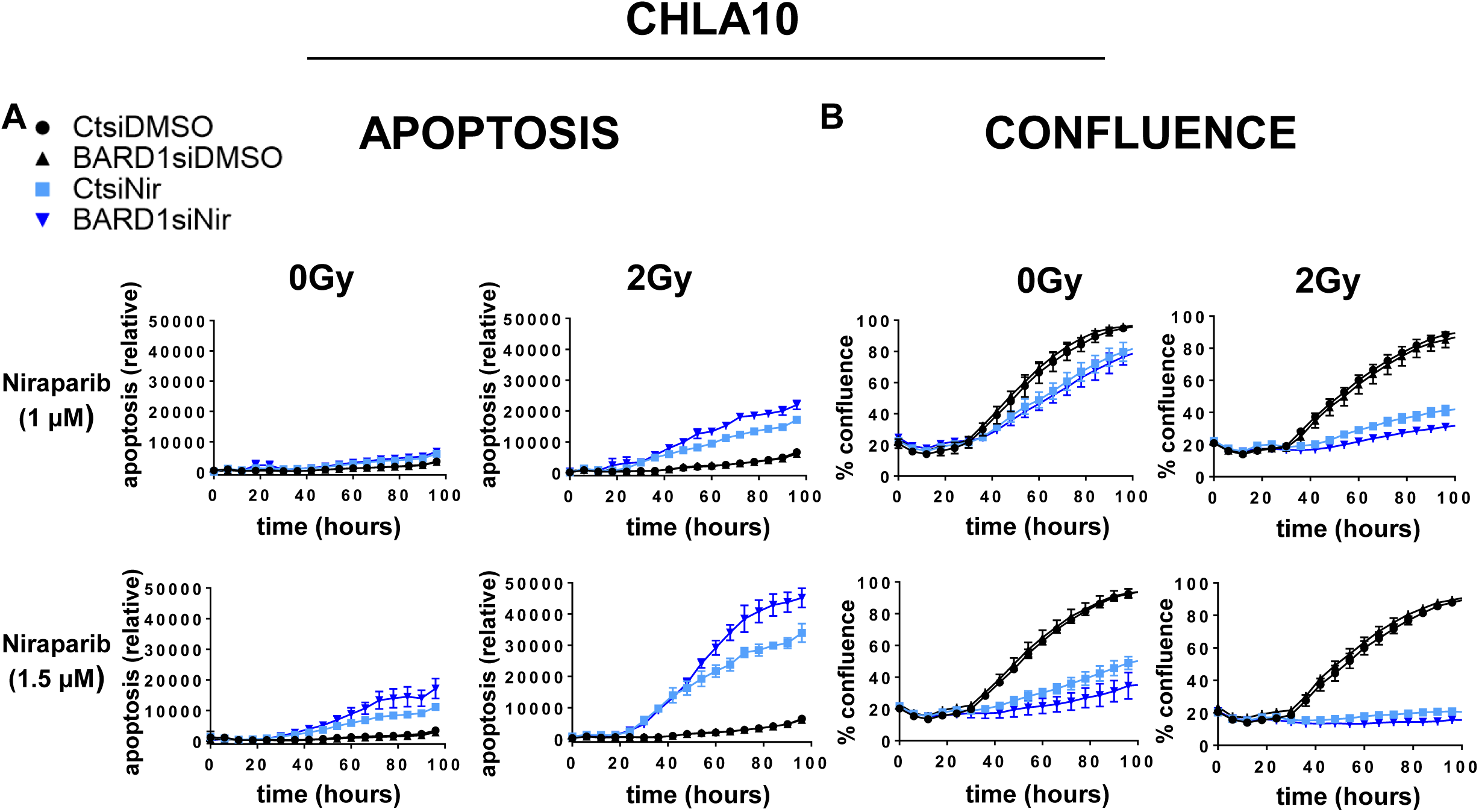
BARD1 loss enhances Ewing sarcoma cell apoptosis in response to niraparib plus radiation in CHLA10 Ewing sarcoma cells. **A, B**, CHLA10 cells were treated with control (Ctsi) or BARD1 (BARD1si) siRNA, Nir (at doses indicated) versus DMSO control, and either 0 or 2 Gy radiation and monitored via IncuCyte apoptosis assay **(A)** or confluence assay **(B)**. For these experiments, cells were seeded in the presence of Nir and radiation was performed at 12-15 hours. Relative apoptosis (caspase 3/7 activity) is calculated as green florescence in μM^2^ divided by percent confluence.

## Notes

Financial Support This work was supported by Alex’s Lemonade Stand Foundation Young Investigator and Million Mile Award (KMB), the Children’s Cancer Research Fund Emerging Scientist Award (KMB), Hyundai Hope on Wheels Young Investigator Award (KMB), The Morden Foundation (KMB), St. Baldrick’s Fellowship Award (LMM) and K12HD052892-13 (LMM, PI Dermody K12HD052892-13). The University of Pittsburgh holds a Physician-Scientist Institutional Award from the Burroughs Welcome Fund (JDD). Research reported here has been supported by the National Institutes of Health under their award number T32HD071834 “Research Training Program for Pediatric Subspecialty Fellows” PI: Terence S. Dermody, M.D.

### Competing Interest Statement

The authors have declared no competing interest.

